# A novel dimeric active site and regulation mechanism revealed by the crystal structure of iPLA_2_β

**DOI:** 10.1101/196097

**Authors:** Konstantin R. Malley, Olga Koroleva, Ian Miller, Ruslan Sanishvili, Christopher M. Jenkins, Richard W. Gross, Sergey Korolev

**Affiliations:** Edward A. Doisy Department of Biochemistry and Molecular Biology, Saint Louis University School of Medicine, St. Louis, MO 63104, USA; GM/CA@APS, Advanced Photon Source, Argonne National Laboratory, Argonne, IL, USA; Division of Bioorganic Chemistry & Molecular Pharmacology, Department of Medicine, Washington University School of Medicine, 660 South Euclid Avenue, Campus Box 8020, Saint Louis, Missouri 63110, USA; Department of Developmental Biology, Department of Medicine, Washington University School of Medicine, Saint Louis, Missouri 63110, USA; Center for Cardiovascular Research, Department of Medicine, Washington University School of Medicine, Saint Louis, Missouri 63110,USA; Department of Chemistry, Washington University, Saint Louis, Missouri 63130 USA

## Abstract

Calcium-independent phospholipase A_2_β (iPLA_2_β) regulates several physiological processes including inflammation, calcium homeostasis and apoptosis. It is linked genetically to neurodegenerative disorders including Parkinson’s disease. Despite its known enzymatic activity, the mechanisms underlying pathologic phenotypes remain unknown. Here, we present the first crystal structure of iPLA_2_β that significantly revises existing mechanistic models. The catalytic domains form a tight dimer. The ankyrin repeat domains wrap around the catalytic domains in an outwardly flared orientation, poised to interact with membrane proteins. The closely integrated active sites are positioned for cooperative activation and internal transacylation. A single calmodulin binds and allosterically inhibits both catalytic domains. These unique structural features identify the molecular interactions that can regulate iPLA_2_β activity and its cellular localization, which can be targeted to identify novel inhibitors for therapeutic purposes. The structure provides a well-defined framework to investigate the role of neurodegenerative mutations and the function of iPLA_2_β in the brain.

## Introduction

Calcium-independent phospholipase A_2_β (iPLA_2_β, also known as PLA2G6A or PNPLA9) hydrolyses membrane phospholipids to produce potent lipid second messengers^1,2^. Due to its emerging role in neurodegeneration and a strong genetic link to Parkinson’s disease (PD)^3-11^, the gene coding for iPLA_2_β was designated as PARK14. Originally isolated from myocardial tissue as an activity stimulated during ischemia^12,13^, the enzyme displays several specific features including calcium-independent activity, a preference for plasmalogen phospholipids with arachidonate at the *sn*-2 position, an interaction with ATP^14,15^ and inhibition by calmodulin (CaM) in the presence of Ca^2+^ ^16^. It was also isolated from macrophages, where it was thought to act as a housekeeping enzyme, maintaining the homeostasis of the lipid membrane^14^. Subsequent studies using the mechanism-based inhibitor bromoenol lactone (BEL) revealed involvement of the enzyme in 1) agonist-induced arachidonic acid release^17^; 2) insulin secretion^18^; 3) vascular constriction/relaxation by Ca^2+^ signaling through store-operated calcium-entry (SOCE) ^19,20^; 4) cellular proliferation and migration^21,22^; and 5) autophagy^23,24^. Alterations in iPLA_2_β function have demonstrated its role in multiple human pathologies including cardiovascular disease^1,25-27^, cancer^28-31^, diabetes^32,33^, muscular dystrophy^34^, nonalcoholic steatohepatitis^35^ and antiviral responses^36^. Correspondingly, novel inhibitors of iPLA_2_β have been sought for therapeutic applications^37,38^. Highly selective mechanism-based fluoroketone inhibitors were designed^37,39,40^ and successfully applied in mouse models of diabetes^41^ and multiple sclerosis^42^. Recently, numerous mutations have been discovered in patients with neurodegenerative disorders such as infantile neuroaxonal dystrophy (INAD)^43-45^ and PD^3-11^. The protein was also found in Lewy bodies and its function was connected to idiopathic PD^24,46^.

More than half of the iPLA_2_β sequence is comprised of putative protein interaction domains and motifs (Figs. 1a, S1). The sequence can be divided into three parts: the N-terminal domain, the ankyrin repeat (AR) domain (ANK) and the catalytic domain (CAT)^47^. Hydrolysis by iPLA_2_β is executed by a Ser-Asp catalytic dyad in close spatial proximity to a glycine-rich motif. The CAT domain is homologous to patatin, a ubiquitous plant lipase^48^. The AR is a 33-residue motif consisting of a helix-turn-helix structure followed by a hairpin-like loop forming a conserved L-shaped structure. ARs are found in thousands of proteins and have evolved as a highly specific protein recognition structural scaffold^49,50^. In different proteins, 4 to 24 ARs can be stacked side-by-side forming elongated linear structures. Five conserved amino acids form a hydrophobic core holding the helical repeats together. The remaining amino acids are variable, but the 3D structure of the AR is highly conserved.

**Figure 1.**
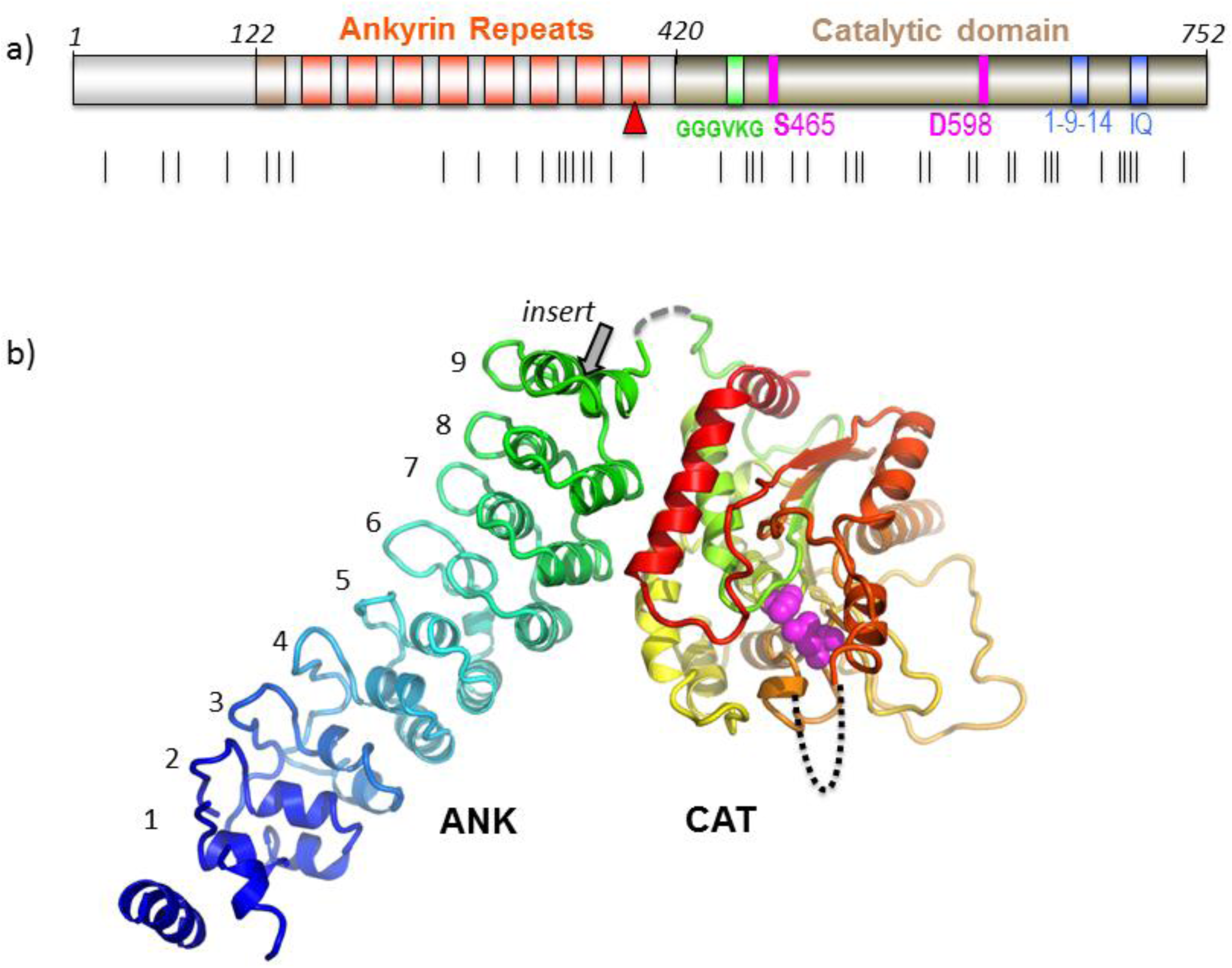
**a)** Domain composition of iPLA_2_β. ARs are shown in orange with the novel AR_1_ in dark orange, catalytic residues are in magenta, poly-Gly region is in green and putative CaM-binding motifs in blue. Black lines underneath mark INAD and PD mutations. **b)** Cartoon representation of the iPLA_2_β monomer color-coded in a rainbow scheme with the N-terminus in blue and the C-terminus in red. The catalytic dyad is shown by magenta spheres. The location of the unstructured loop between ANK and CAT domains is indicated by dashed grey line and of the disordered membrane-interacting loop by black dotted line. The position of the proline-rich insert in the long variant is shown with an arrow.

**Figure 2.**
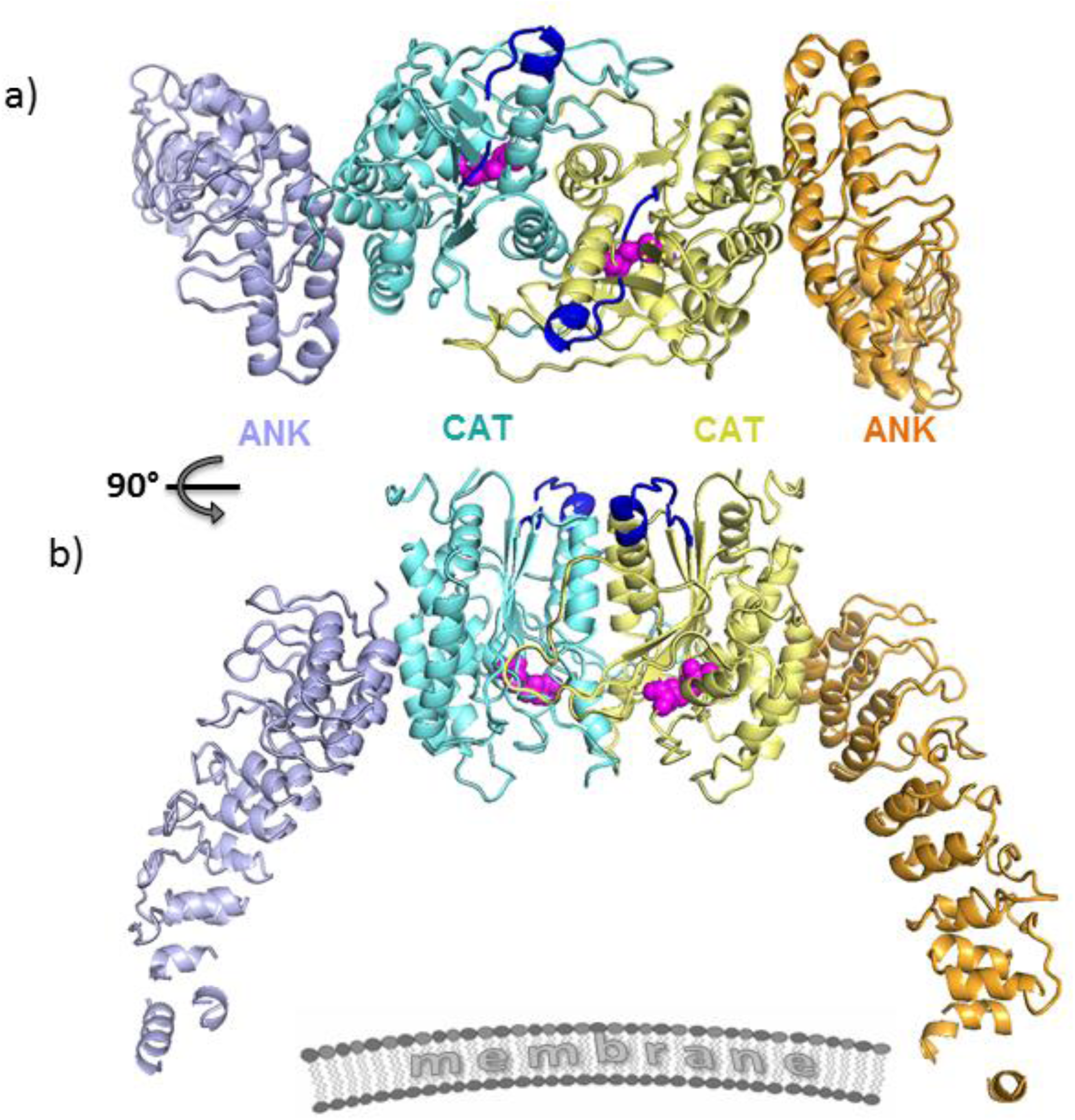
Configuration of the iPLA_2_β dimer in crystal structure. **a**) The CAT and ANK domains of a dimer are shown in cyan and light navy, respectively, in monomer A and in yellow and orange in monomer B. Putative CaM-binding 1-9-14 motifs in both monomers are shown in dark blue. Catalytic dyads are shown by magenta spheres. **b)** Same dimer rotated by 90° around horizontal axis. The schematic drawing of membrane illustrates the orientation of the membrane-binding surface of iPLA_2_β.

The cellular localization of iPLA_2_β is tissue-specific and dynamic^13,20,51^. Different variants of iPLA_2_β are associated with the plasma membrane, mitochondria, endoplasmic reticulum and the nuclear envelope. iPLA_2_β lacks trans-membrane domains, but is enriched in putative protein-interaction motifs. Those include several proline-rich loops and the extended ANK domain with 7 or 8 ARs capable of interacting with multiple cognate receptor proteins^49,50,52^. However, relatively little is known about iPLA_2_β protein interaction mechanism. It binds calmodulin kinase (CaMKIIβ) in pancreatic islet β-cells^53^ and the ER chaperone protein calnexin (Cnx)^54^. The functional significance and mechanisms of both interactions remains unknown. Pull-down of proteins isolated from β-cells under mild detergent treatment revealed a number of other proteins from different cellular compartments, including transmembrane proteins^54^. iPLA_2_β was also found in the Arf1 interactome, which regulates cell morphology^55^. Understanding the mechanisms of the diverse iPLA_2_β functions requires knowledge of its spatial and temporal localization, which are most likely guided by poorly understood protein-protein interactions. Overall, structural studies are currently limited to identification of the putative calmodulin binding sites^56^, molecular modeling, and mapping of the membrane-interaction loop using hydrogen/deuterium exchange mass spectrometry^57-59^.

Here, we present the first crystal structure of a mammalian iPLA_2_β, which revises previous structural models and reveals several unexpected features critical for regulation of its catalytic activity and localization in cells. The protein forms a stable dimer mediated by CAT domains, with the active sites poised to interact cooperatively, facilitating transacylation and, potentially, other acyl transfer reactions. The structure suggests an allosteric mechanism of the inhibition, where a single CaM molecule interacts with two CAT domains altering the conformation of dimerization interface and active sites. Surprisingly, ANK domains in the crystal structure are oriented toward the membrane interaction interface and are ideally positioned to interact with membrane proteins. Structural data suggest a novel ATP-binding site in the AR and the role of ATP in regulating protein activity.

## Results

### Structure of iPLA_2_β

The structure of the short variant of iPLA_2_β (SH-iPLA_2_β, 752 amino acids) was solved by a combination of selenomethionine single-wavelength anomalous diffraction (SAD) with molecular replacement (MR) using two different protein models. Those include the patatin^48^, with 32% sequence identity to the CAT domain, and four ARs of an ankyrin-R protein^60^ (Fig. S1). Five additional ARs and several loop regions in CAT were modeled into the electron density map. The sequence assignment was guided by position of 51 selenium peaks and the structure was refined using 3.95 Å resolution data (Table S1, Fig. S2). Residues 1-80, 95-103, 113-117, 129-145, 405-408, 630-631 and 652-670 were omitted from the final model. Regions 81-94, 104-112 and 409-416 were modeled as alanines. The short variant lacks a proline-rich loop in the last AR (Fig. 1) and sequence numbering in the paper corresponds to sequence of the SH-iPLA_2_β. The structure of the monomer is shown in Figure 1b.

The core secondary structure elements of the CAT domain are similar to that of patatin with root-mean-square deviation (rmsd) of 3.1 Å for 186 Cα atoms (Fig. S3a). Consequently, the fold of the CAT domain also resembles that of cPLA_2_α catalytic domain^61^, but to a significantly lesser extent. The active site is localized inside the globular domain as in the patatin structure. However, in iPLA_2_β, the catalytic residues are more solvent-accessible than in patatin (Fig. S3b). In the latter, the active site is connected to the surface through two narrow channels (Fig. S3c), insufficient for phospholipid binding without significant conformational changes. By contrast, in iPLA_2_β, the active site cavity is wide open and can accommodate phospholipids.

The periphery and loop regions differ significantly from those in the patatin structure, with two unique extended proline-rich loops in iPLA_2_β. A long C-terminal α-helix (α7 in patatin^62^) is kinked in the iPLA_2_β structure and participates in dimerization (described below).

### Conformation of the ANK domain

The electron density map reveals nine ARs in the structure of SH-iPLA_2_β, instead of the previously predicted eight. AR_1_ is formed by residues 120-147 with a less conserved AR signature sequence motif (Fig. S1). The outer helix of AR_1_ is poorly ordered and was omitted from the current model. The C-terminal AR_9_ is formed by residues 376-402. Gln396, which is substituted by the 54-residue proline-rich insert in the long variant (L-iPLA_2_β), locates in the short loop connecting two helixes of AR_9_ (grey arrow in Fig. 1b). The orientation of the entire ANK domain is completely unexpected (Figs. 1b, 2b). It is attached to the CAT domain at the side opposite to the membrane-binding surface and was expected to form an extended structure oriented away from the membrane to participate in oligomerization^63^. In the crystal structure, it wraps around the CAT domain towards the predicted membrane interfacial surface. This is achieved by the extended conformation of an eighteen amino acid-long connecting loop, illustrated in Figure S6d. Part of the linker is unresolved due to poor electron density, however, the assignment of the ANK and CAT domains to the same molecule is unambiguous in the crystal packing. The outer helices of AR_7_ and AR_8_ form an extensive hydrophobic interface with CAT. AR_9_ partially contributes to this interface as well.

### ANK interaction with ATP

iPLA_2_β is the only known phospholipase that interacts with ATP^12^. The glycine-rich motif was initially proposed as an ATP-binding site^64^. However, this motif is highly conserved through patatin-like phospholipases, where it forms part of the active site. It is also a common element of α/β hydrolases, where it functions as an oxyanion hole coordinating charge distribution during catalysis^65^. To identify the location of ATP binding in iPLA_2_β, we soaked protein crystals with 2’MeSe-ATP and collected 4.6 Å anomalous data. A single anomalous peak was consistently found near Trp293 of AR_6_ (Fig. S6a). An electron density near this residue was also found in the 2Fo-Fc map calculated from the SeMet crystal (Fig. S6b), confirming a potential interaction with ATP at this location. The density was less pronounced in the native crystal, and since the low resolution of the Se-Met and Se-ATP data did not permit unambiguous modeling of ATP, we did not include it in the final refinement. Importantly, AR_6_ (residues 282-308) adopts an unusual conformation. One of its helices is two amino acids shorter than a conventional AR helix. There is a kink of the entire ANK domain at this position, as compared to ankyrin-R. Potential binding of ATP at this location, where an elongated ANK domain structure is disrupted by the short α-helix of AR_6_, suggests the importance of ATP to the regulation of ANK domain conformation and thermodynamic stability.

To our knowledge, the interaction of ATP with ARs has been reported only once in the literature. TRPV1 binds ATP within the positively charged inner concave surface of three ARs^66^. In iPLA_2_β, AR_6_ and AR_7_ also possess several basic residues in a corresponding surface area. However, the position of the anomalous peak next to Trp293 suggests a potential stacking interaction. Interestingly, it was shown that both purine nucleotides, ATP and GTP, have similar effects on iPLA_2_β activity^15,67^.

### CAT-mediated dimerization of iPLA_2_β

The crystallographic asymmetric unit is formed by a dimer of iPLA_2_β. In contrast with the original hypothesis of ANK-mediated oligomerization^63^, this dimer is formed through CAT domains. The ANK domains are oriented outwards in opposite directions, forming a ∼150 Å-long elongated structure. Analytical ultracentrifugation (AUC) experiments support the existence of an elongated dimeric structure in solution (Fig. 4a). The experimental MW was 152 kDa, corresponding to a dimer with a theoretical MW of 170 kDa. The friction ratio of 1.9 (compared to 1.4 for BSA) corresponds to a significant deviation from a globular shape.The isolated ANK domains did not oligomerize in an AUC experiments (Fig. S4b).

CAT domains interact through an extended, largely hydrophobic dimerization interface with a contact area of ∼2800 Å^2^ (Fig. 3) formed by three α-helices, several loops, including the long loop 554-570, and a part of the central β-sheet. Such an extensive interaction supports a stable dimer. Correspondingly, iPLA_2_β dimerizes even at nanomolar concentrations (Fig. S4a). To probe the dimerization interface, we substituted Trp695 with Glu (W695E). Trp695 forms extensive hydrophobic interactions with the opposite monomer, including a stacking interaction with its counterpart (Fig. S6c). The mutant is a monomer in solution and is inactive (Fig. 4A, S4g). The W695A mutant exists in equilibrium with both monomeric and dimeric peaks and is catalytically active (data not shown).

**Figure 3.**
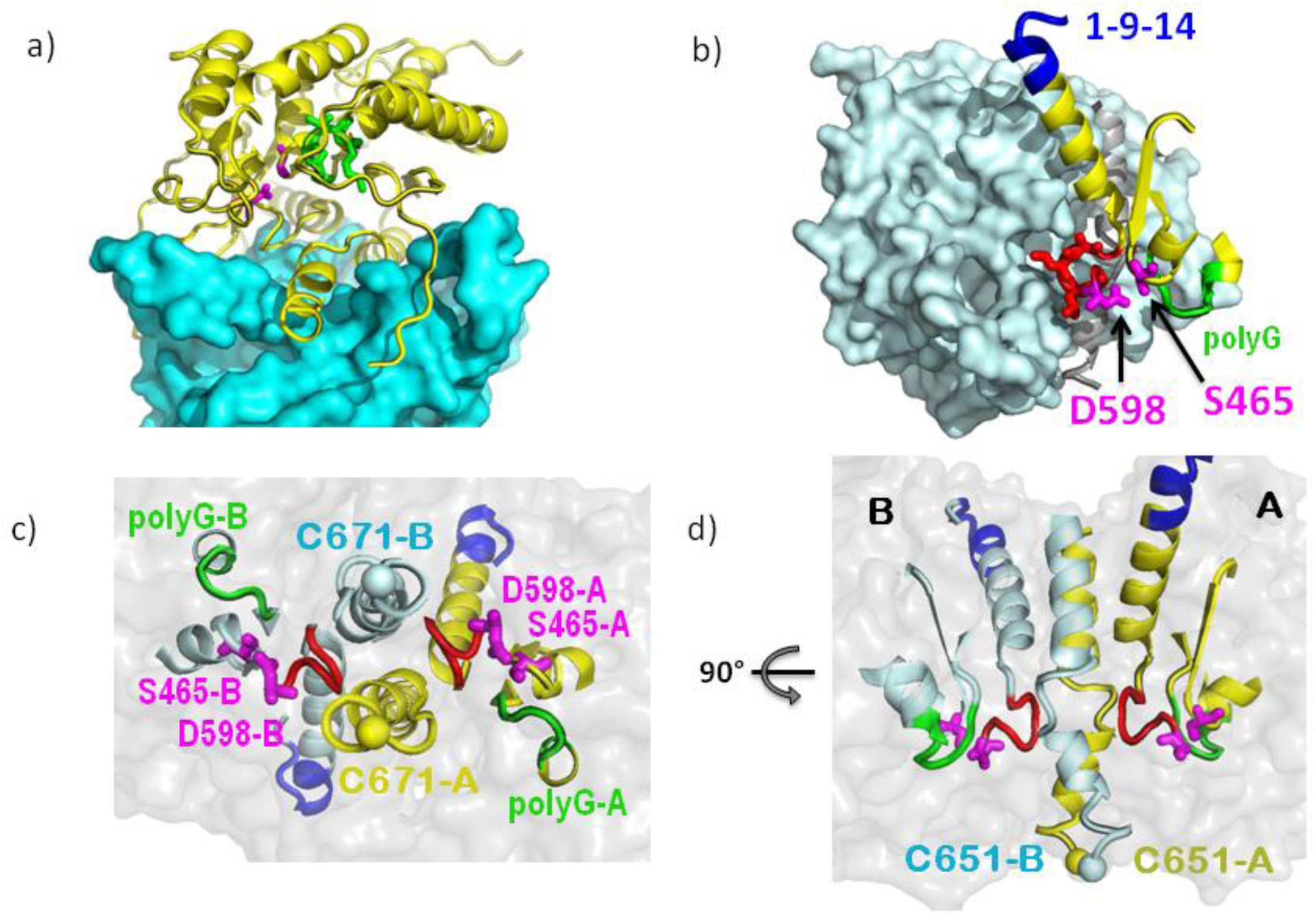
Extensive interactions of CAT domains and integrated active sites. **a)** Interaction of the CAT domain of molecule B (CAT-B), shown as cyan surface, with the CAT domain of molecule A (CAT-A), shown as yellow cartoon with highlighted catalytic dyad residues (magenta sticks) and the oxyanion hole (green). **b)** The proximity of the active site to the dimerization interface is illustrated with surface representation of CAT-B (light cyan) and structural elements of the CAT-A active site shown as yellow cartoon, along with the Ser-Asp dyad of CAT-A (magenta stick representation), the oxyanion hole formed by poly-Gly loop (green), and the π-helix (red) which contains the catalytic Asp. The structured fragment of 1-9-14 motif is shown in blue. **c)** The view from the membrane binding surface of the active sites of a dimer with secondary-structure elements and the individual residues color-coded as in b) for molecule A and by light cyan for molecule B. A transparent surface of the dimer is shown in grey. C671 residues of the dimer are represented by yellow and light cyan spheres. These cysteines were previously reported to be acylated in the presence of acyl-CoA and are located on the membrane side of the protein surface. **d)** Side view of the same structural elements in orientation orthogonal to that in c), illustrating the distance of catalytic dyad residues from the membrane interacting surface and the location of Cys671 at this surface as well as near the dimerization interface.

**Figure 4.**
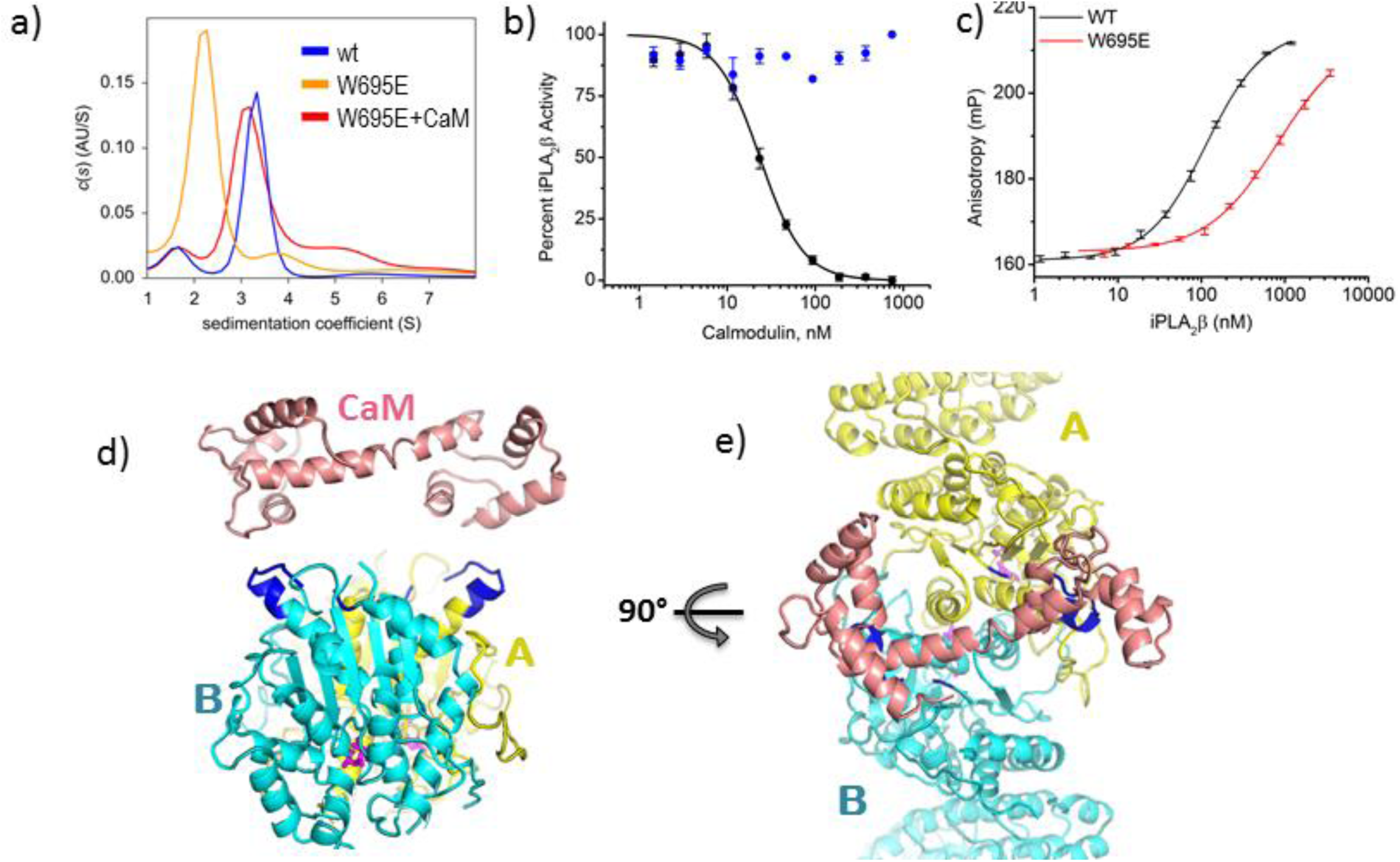
iPLA_2_β dimerization and mechanism of CaM inhibition. **a)** Sedimentation velocity s(w) distributions of wild type iPLA_2_β shown in blue (peak MW: 152 kDa, theoretical MW: 170 kDa), of the W695E mutant in orange (83 kDa), and of W695E in the presence of CaM/Ca^2+^ in red (138 kDa). Lower MW can be due to unresolved contribution of minor fraction of monomeric species in a latter case. **b)** Inhibition of iPLA_2_β enzymatic activity by CaM in the presence (black) and absence (blue) of Ca^2+^. **c)** Interaction of FAM-CaM/Ca^2+^ with iPLA_2_β as measured by fluorescence anisotropy upon titration by wild-type iPLA_2_β (black) and the W695E mutant (red). Error bars represent average ± s.e.m of triplicate experiments, which were performed at least twice independently. **d)** and **e),** two orthogonal views of the iPLA_2_β dimer with monomers color coded in cyan and yellow. CaM in the conformation as reported in the 3SJQ PDB structure is placed next to 1-9-14 motifs (highlighted in blue) to illustrate the possibility of CaM interaction with two 1-9-14 motifs of a dimer.

The monomers are related by a two-fold axis rotational symmetry. Two active centers and the predicted membrane-binding loops^57^ are oriented in the same direction (Fig. 3d). Importantly, the active sites are in the immediate vicinity of the dimerization interface and in close spatial proximity to each other (Fig. 3). The catalytic Asp598 is at the beginning of a π-helical loop (599-603) and two leucines of this loop form contacts with the long α-helix (604-624) of the opposite monomer. This arrangement suggests a strong allosteric association between the two active sites and dependence of the catalytic activity on the dimer conformation.

### Calmodulin binding mechanism

Calmodulin inhibits iPLA_2_β enzymatic activity in the presence of calcium. It was proposed to tightly interact with iPLA_2_β even at low calcium concentrations^68^ and to be displaced by active mechanisms, such as covalent modification of the active site by acyl-CoA^69^ or by interaction with a calcium influx factor released from the ER during calcium depletion^70^. Two putative CaM-binding peptides containing the canonical IQ and 1-9-14 motifs were previously isolated by tryptic footprinting and affinity chromatography using CaM-agarose^56^. We measured the K_i_ of iPLA_2_β inhibition by CaM using a fluorogenic activity assay with Pyrene-PC fluorescent phospholipid liposomes (Fig. S5a-e). The results revealed a tight calcium-dependent interaction with CaM with a K_i_ of 23 ± 1.5 nM (Fig. 4b) and a Hill coefficient n = 2.2 ± 0.2, indicating potential cooperativity. Next, we measured the direct interaction of CaM with iPLA_2_β using fluorescent polarization with fluorescein (FAM)-labeled CaM (Fig. 4c). The dissociation constant of the interaction of CaM with iPLA_2_β (K_d_) of 112 ± 5 nM was higher than the K_i_ measured with unmodified CaM; however, it corresponds to the K_i_ of FAM-CaM (Fig. S5f). No cooperativity was observed in the direct binding experiment.

Remarkably, the interaction of FAM-CaM with the monomeric W695E mutant was at least an order of magnitude weaker, with a K_d_>1400 nM (Fig. 3c), suggesting that iPLA_2_β dimerization is crucial for CaM binding. The interaction of CaM with synthetic isolated FAM-labeled peptides corresponding to 1-9-14 and IQ motifs was even weaker. Affinity towards the 1-9-14 motif (K_d_ = 2500 ± 400 nM) was comparable to that of the monomeric W695E mutant. Binding of the IQ motif was even weaker (K_d_=5900 ± 800 nM) (Fig. S5h).

Finally, an excess of CaM enabled W695E dimerization in sedimentation velocity AUC experiments (Fig. 3a). These data strongly support the model where a single CaM molecule interacts with an iPLA_2_β dimer and explain potential cooperativity in the inhibition assay. Furthermore, the two 1-9-14 motifs are located on the same side of the dimer and are ∼30 Å apart from each other (Figs. 4d,e). In the structure of the small conductance potassium channel complex with CaM (PDBID: 3SIQ)^71^, a single CaM molecule in an extended conformation interacts with the channel dimer and the distance between CaM-binding helixes is also 30 Å. In Figures 4d and 4e, CaM from the 3SIQ complex is placed next to an iPLA_2_β dimer to illustrate comparable distances. At the same time, the conformation of the IQ motif in tertiary structure makes it as unlikely target of CaM binding. This motif overlaps with a β-strand of the conserved structural core of the molecule and is inaccessible for binding without protein unfolding. Moreover, mutation of the most conserved hydrophobic Ile to a charged Asp (I701D) in the IQ motif did not affect iPLA_2_β inhibition by CaM (data not shown). Together, results from solution studies and the conformation of potential CaM-binding sites in the iPLA_2_β dimer suggest that one CaM molecule interacts with two monomers of the iPLA_2_β dimer, most likely through the 1-9-14 motifs.

### Discussion

The first crystal structure of iPLA_2_β has revealed several unexpected features underlying its enzymatic activity, mechanisms of regulation and structural domains potentially involved in tissue-specific localization. Previous computer modeling studies used the patatin structure and proposed an interfacial activation mechanism whereby interaction with membrane leads to opening of a closed active site^37^. In the iPLA_2_β crystal structure, the active site adopts an open conformation in the absence of membrane interaction (Fig. S3b). Both active sites of the dimer are wide open and provide sufficient space for phospholipids to access the catalytic centers. This is in contrast to patatin, where only two narrow channels connect the catalytic dyad with the solvent exposed surface, and conformational changes are required for substrate to access the active site (Fig. S3c). An open conformation of the active site explains the ability of iPLA_2_β to efficiently hydrolyze monomeric substrates^16^ and the lack of a strong interfacial activation such as observed with cPLA_2_, where membrane binding increases activity by several orders of magnitude^72^.

### Dimeric active site

The dimer is formed by CAT domains tightly interacting through an extensive interface, while ANK domains are oriented outwards from the catalytic core. The existence of the dimer in solution was confirmed by quantitative sedimentation velocity and cross-linking experiments. This configuration was verified by mutagenesis of the observed dimerization interface and a lack of oligomerization of isolated ANK domains. The elongated shape of the dimer contributes to an overestimation of the previously reported oligomeric state in gel filtration analysis due to faster migration of elongated molecules through the size-exclusion matrix. A remote iPLA_2_β homolog from *C. elegans* also forms a dimer in solution^24^.

The catalytic centers are in immediate proximity to the dimerization interface and the activity is likely to depend on the conformation of the dimer. Disruption of the dimer by the W695E mutation yields an inactive enzyme. The active sites are also in close proximity to each other and allosterically connected. Concerted activation of closely integrated active sites should promote rapid responses upon stimulation by ligands, rendering the enzyme an efficient sensor of external perturbations.

Close proximity of active sites provides a plausible explanation of the previously reported activation mechanism through autoacylation of Cys651. The reaction occurs in the presence of oleoyl-CoA and the modified enzyme is active even in the presence of CaM/Ca^2+^ ^69^. Cys651 is located at the entrance to the active site at the base of the membrane-binding loop as well as at the dimerization interface (Fig. 3d). Covalent attachment of a long fatty acid chain at this position should increase protein affinity to membrane and can alter the conformation of a CaM-bound dimer. The close proximity of two active sites provides an explanation for this autoacylation phenomenon important for the mechanism of enzyme activation in the heart during ischemia.

An intimate allosteric connection of active sites and the dimerization interface provides a plausible mechanism for inhibition by CaM. Indeed, solution studies and location of the putative CaM-binding site strongly suggest that a single CaM binds two molecules of the dimer. We hypothesize that such interactions will lead to conformational changes in the dimerization interface and alter conformation of both active sites.

A hypothetical model of two potential states of iPLA_2_β with CaM-bound inactive and CaM-free active dimers is illustrated in Figure 5. In both states, the enzyme is a dimer. The conformation of the dimerization interface differs in the two states depending on interaction of CaM with the 1-9-14 motif. Allosterically, CaM binding stabilizes a closed conformation of the active sites, which remain open in the absence of CaM. The positions of ATP and of acyl modification are shown in the active form. However, the exact mechanism of activation through autoacylation and the effect of ATP-binding on protein activity remain to be further investigated. ANK domains are likely to move out of the conformation observed in crystal structure upon approaching the membrane. In crystals the dimer is shaped as an arch standing on legs formed by the ANK domains (Fig. 2b), with the CAT domains at the top and with their active sites facing downward. The inner radius of the arch is ∼80-100 Å. Therefore, in this conformation the ANK domains can prevent the membrane surfaces of larger radii from accessing the catalytic domains. However, the non-specific hydrophobic interactions permit rotational flexibility of interacting domains. Therefore, ANK domains can rotate out of this inhibitory position, while maintaining hydrophobic contacts with the CAT domain. In fact, the relative orientation of the ANK and CAT domains is slightly different between the two monomers of the same crystal. Upon superposition of CAT domains of two monomers, the resulted orientation of ANK domains differs with N-terminal ends shifted by ∼12 Å (Fig. S6e). Similar variation of the ANK domain orientation is a major source of non-isomorphism between different crystals, as observed between native and SeMet crystal forms (data not shown).

**Figure 5.**
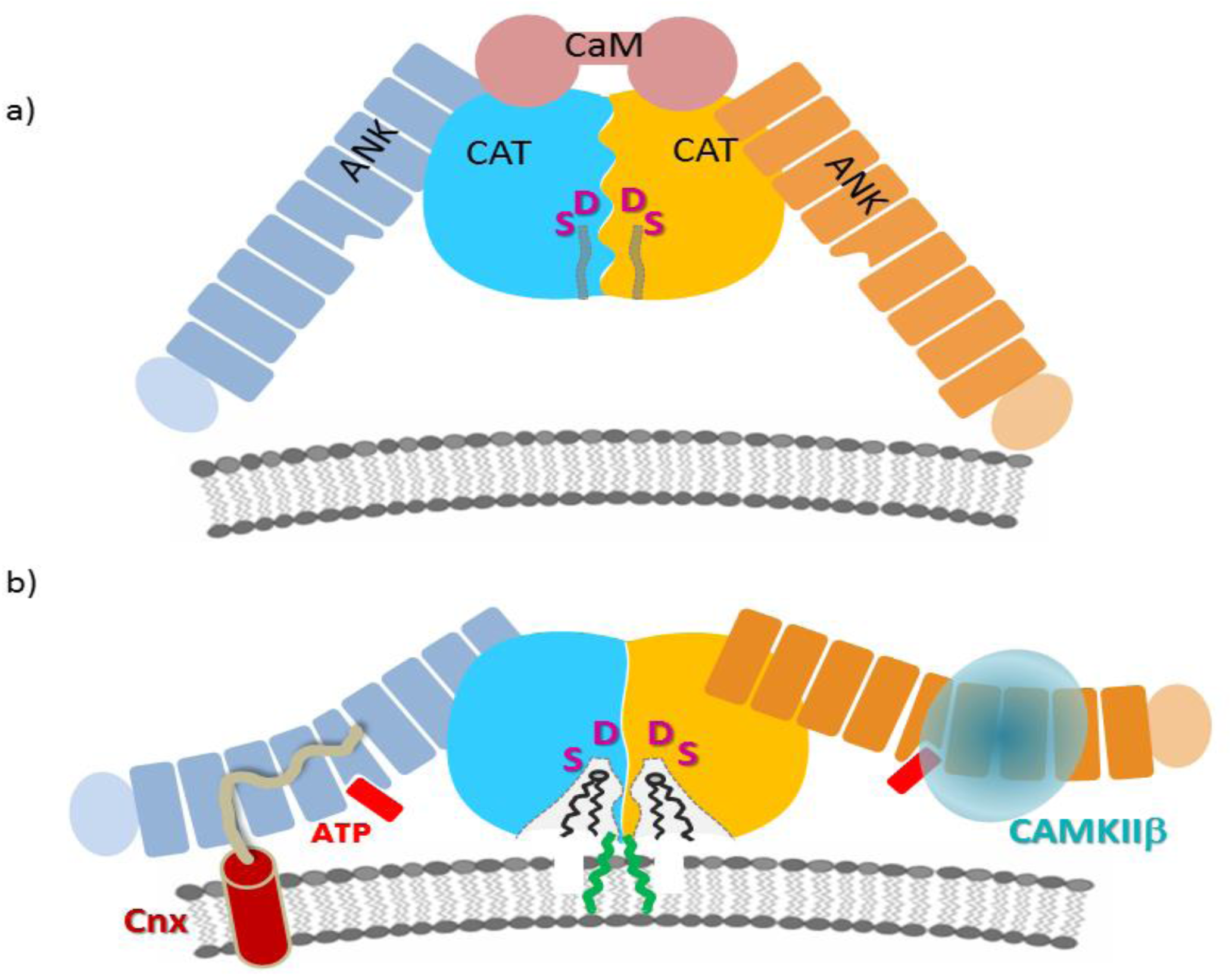
The proposed mechanism of iPLA_2_β regulation and macromolecular interactions. **a)** <u>Schematic representation of iPLA_2_β dimer in a hypothetical inhibited state bound to CaM</u>. CAT domains are shown in blue and yellow, ANK domains in navy and orange, and a single CaM molecule is represented by two connected circles in pink. Active site cavities are represented by narrow channels (grey lines) leading from the solvent-exposed surface to the Ser/Asp catalytic dyad depicted by magenta. **b**) <u>An active conformation of the dimer</u>. CaM dissociation leads to the opening of the active sites. ANK domains are available for interactions with protein partners as illustrated for CAMKIIβ (light cyan transparent sphere), known to interact with ANK domain, and with transmembrane Cnx (shown as transmembrane helix with the C-terminal cytosolic peptide in pale yellow), which could recruit iPLA_2_β to the membrane. The Cnx-binding site of iPLA_2_β is not known and the hypothetical interaction with ANK domain is based on similar interaction of AnkB and sodium channel peptide. ATP binding (red) in the middle of the ANK domain could trigger additional conformational changes of the AR. Acylation of C671 by oleoyl-coA (green) can facilitate interaction with the membrane and/or opening of active site channels. Other conformational states are feasible as well, such as CaM-bound inhibited protein at the membrane or an open conformation of active sites in CaM-free form in cytosol, corresponding to the crystallized form.

**Figure 6.**
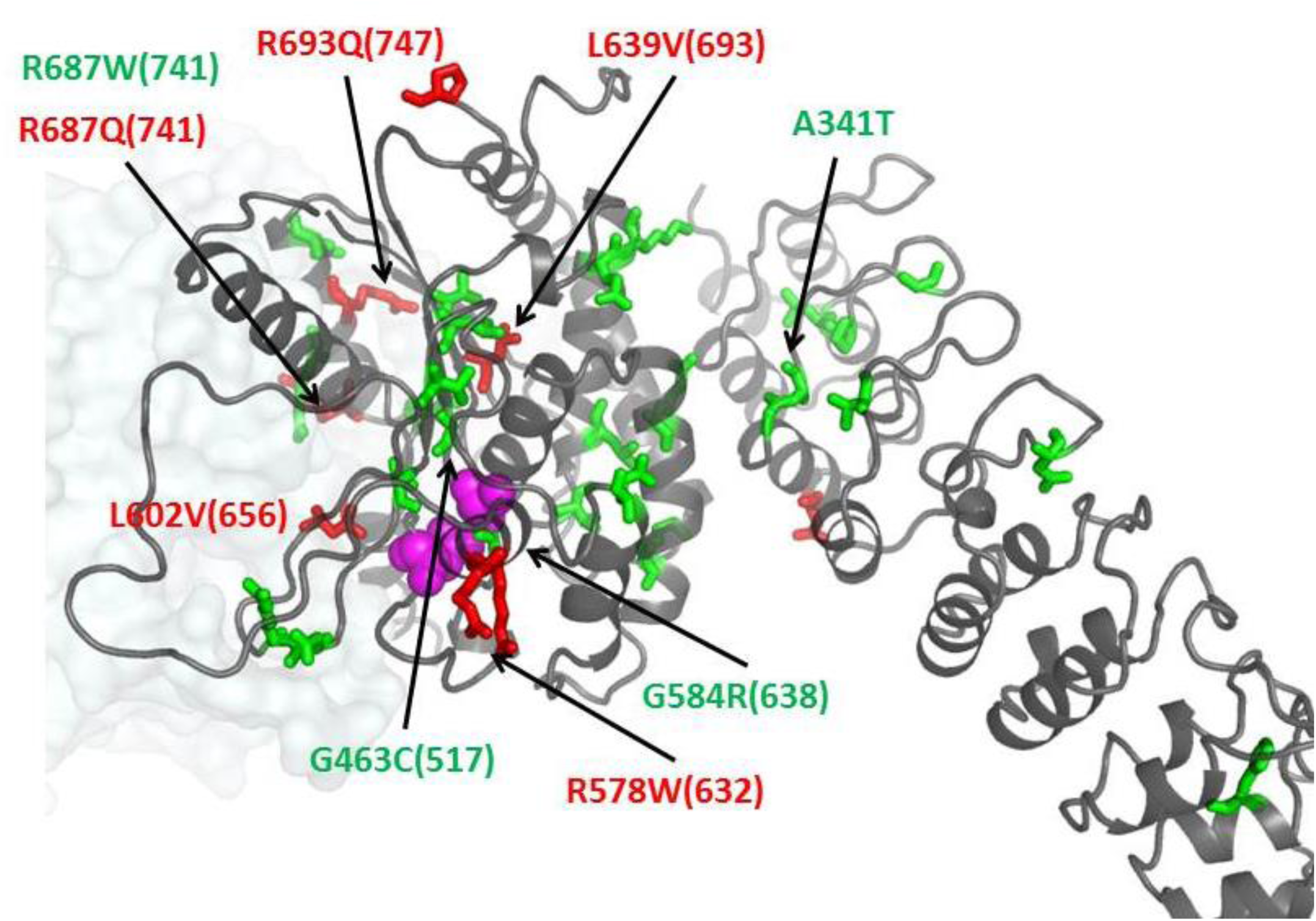
Position of selected INAD and PD mutations. Residues mutated in INAD and PD patients are shown as green and red sticks, respectively, on the cartoon of the iPLA_2_β monomer. Position of the second monomer is shown as pale transparent surface. Active site Ser/Asp residues are shown by magenta spheres. Mutations previously tested for enzymatic activity are labeled. Numbers in parentheses correspond to the human long variant (L-iPLA_2_β) sequence.

The model also illustrates a hypothetical interaction of ANK domains with cytosolic CAMKIIβ and with cytosolic C-terminus of transmembrane calnexin, discussed below.

### Novel structure suggests a role for ANK domains in membrane localization

The iPLA_2_β field currently lacks a coherent explanation of how it localizes to the membrane versus cytosol in different cell types and tissues. Elongated ARs form highly specific docking surfaces for different protein partners and 4-8 ARs are capable of binding several proteins^73^. The finding that the ANK domains are oriented towards the membrane-facing side of iPLA_2_β suggests that ARs represent the most logical site of putative interaction with membrane proteins. Indeed, specific protein recognition is the only known function of ARs. The canonical ARs tether cytoskeletal proteins to ion channels^60^.

The interaction of CAMKIIβ with the N-terminal part of iPLA_2_β has been reported. The function of such interaction remains poorly understood. It can either modulate activity of each protein or mediate recruitment of iPLA_2_β to membrane since CaMKIIβ form complexes with actin, an L-type Ca^2+^ channel regulatory protein a1C, and synaptic proteins like NMDA receptors, densin-180 (a trans-synaptic protein), and α-actinin (an actin-binding protein). In case of strong interaction, CaMKIIβ may also play a role of iPLA_2_β storage pool due to high abundance of protein in neurons (2% of total protein in hippocampus).

The Cnx-binding domain of iPLA_2_β remains unknown. This interaction is particularly interesting in light of growing number of data implicating iPLA_2_β function in ER stress response in β-cells and neurons^24,74,75^. Cnx is the ER chaperone protein. It consists of the lumenal domain, single transmembrane helix and 90 amino acids long C-terminal cytosolic tail, which can interact with iPLA_2_β. Interestingly, interaction of elongated unstructured peptides was previously reported for the AnkB protein with an autoinhibitory peptide and a peptide of the Nav1.2 voltage-gated sodium channel^76^. Hypothetically, the ANK domain of iPLA_2_β could similarly interact with a portion of Cnx C-terminal peptide.

The proline-rich 54-residue insert in the long variant is predicted to form an unstructured loop protruding away from AR_9_, which can also interact with other proteins. Alternatively, it can disrupt the conformation of AR_9_ and alter orientation of the ANK domain. The hydrophobic interface between ANK and CAT domains and long flexible linker can allow for significant movement of the ANK domain, similarly to those observed in Rep helicase^77^ or prothrombin^78^. Additional studies are required to uncover the mechanism and functionality of alternative conformations of iPLA_2_β.

### Neurodegenerative mutations

are found in all domains and therefore can affect the enzymatic activity, regulation or protein interaction properties of iPLA_2_β. In 2006, INAD was linked to mutations in iPLA_2_β gene (PARK14)^43^, which later was connected to a spectrum of neurodegeneration disorders, correspondingly termed PLAN (recent summary and references in ^79^). Those include infantile neuroaxonal dystrophy (INAD1/NBIA2A), atypical neuroaxonal dystrophy (NAD) and idiopathic neurodegeneration with brain iron accumulation including Karak syndrome (NBIA2B). A different set of mutations was linked to a rapidly progressive young-adult onset dystonia-Parkinsonism (PARK14)^5,7,9,11,80-82^. As shown in Figures 1a and 6, mutations are spread throughout all domains. Several tested PARK14 mutants retain full^24,83^ or partial activity^5^, while several tested INAD mutations lead to catalytically inactive enzyme^83^. An interesting example of sensitive allosteric regulation is Arg 687 (corresponding number in L-iPLA_2_β is 741) located at the dimerization interface, which is mutated to Trp in INAD, leading to an inactive enzyme, and to Gln in PD with the activity retained. While an Arg to Trp mutation can significantly alter conformation of the dimerization interface important for catalytic activity, it is unclear what effect a minor Arg to Gln mutation will have and why it cause a late offset (comparatively to INAD) disease. Surprisingly, the A341T mutation in the ANK domain was found to be inactive^83^. This residue is at the ANK/CAT interface and can affect the interactions and stability of the protein. It should be noted that there are very few enzymatic and biochemical studies of the protein and mutants, mostly limited to semi-quantitative measurements. The new structure will facilitate in-depth analysis of known mutants and their effect on biochemical properties. This will lead to a better understanding of protein function and the mechanism of activity and regulation in numerous cellular pathways and disease states.

The structure should also facilitate ongoing design of small molecule modulators of iPLA_2_β for therapeutic purposes. Combined with the analysis of disease-associated mutations, our results clearly demonstrate the importance of N-terminal and ANK domains as well as of peripheral regions of the CAT domain, such as the dimerization interface, for the catalytic activity and its regulation. Together with further knowledge of iPLA_2_β binding partners, such allosteric regions can be targets for small molecule binding to inhibit either enzymatic functions or signaling in downstream pathways.

### Methods

## Protein purification

The pFastBac vector containing iPLA_2_β cloned from CHO cells with a C-terminal 6XHisTag was used for protein expression as previously described^16^.The CHO iPLA_2_β protein was expressed in Sf9 cells using the Bac-to-Bac system (Invitrogen). Bacmid DNA was transfected into Sf9 cells with Trans-IT transfection reagent (Mirus Bio). After 4 days, the media was collected as the p0 viral stock. This stock was amplified by adding 1 mL of p0 to 100 ml of 2x10^6^ cells/mL for 96 h, creating the p1 viral stock. 25 mL of the amplified p1 was used to infect 500 ml shaker flasks of 2x10^6^ cells/mL for 60 h. The cell pellet was washed with cold PBS and suspended in purification buffer (25 mM HEPES pH 7.5, 20% glycerol, 0.5 M NaCl, 1mM TCEP) containing 50 µg/ml each of leupeptin and aprotinin. The cell suspension was frozen in liquid nitrogen and lysed by thawing and sonication at 50% power, 50% duty cycle 4 times for 2 min each. The lysate was cleared by ultracentrifugation at 35,000 rpm for 1 h. 0.5 M urea and 1 mM TCEP were added to the supernatant and mixed with 5 ml of TALON cobalt resin (Clontech) to bind for 1 h at 4°C. The resin was centrifuged at 1,000 rpm for 1 min to remove the flow-through fraction in batch mode. The resin containing the bound protein was then applied to an empty column, washed sequentially with purification buffer containing 10 mM imidazole (100 mL), 40 mM imidazole (40 mL), and eluted with 15 mL purification buffer containing 250 mM imidazole. iPLA_2_β and all mutants were >98% pure as determined by SDS-PAGE and Coomassie staining.

The CaM expression plasmid was a gift from M. Shea (University of Iowa). CaM and its mutants were expressed and purified as previously described^84^.

### Crystallization

iPLA_2_β was concentrated to 6-8 mg/ml in 10 mM HEPES pH 7.5, 500 mM NaCl, 10% glycerol, 5 mM ATP and 1 mM TCEP. Initial crystallization trials were conducted in sitting-drop plates with a Phoenix robot. iPLA_2_β forms crystals within 24 h in several conditions, and after extensive optimization, two primary conditions were selected: 0.1 M bis-Tris pH 5.5, 10% PEG3350, 0.2 M Na/K tartrate and 0.1 M bis-Tris pH 5.5, 10% PEG3350, 0.2 M sodium acetate. Crystals in sitting drop conditions displayed poor diffraction (5-7Å), high X-ray sensitivity and quick deterioration of diffraction power after few days, even while continuing to grow in size. Alternatively, a higher concentration of protein solution was obtained in the presence of CaM. An equimolar amount of purified CaM was mixed with iPLA_2_β, reduced with 5 mM DTT, and dialyzed in 10 mM HEPES pH7.5, 150 mM NaCl, 10% glycerol, 1 mM CaCl_2_. 2 mM ATP was added and the proteins concentrated to 10-12 mg/ml. However, the crystals obtained from iPLA_2_β in the presence of CaM were identical to those obtained without CaM and SDS-PAGE analysis demonstrated the absence of CaM in the crystals.

Growth of suitable protein crystals (diffracting to better than 4Å resolution) was enabled by the counter-diffusion method in capillaries, originally using the Granada Crystallization Box (Hampton Research)^85^, and, later, using a modified capillary method. This method relies on precipitant diffusing through an agarose plug and mixing with the protein solution pre-filled into a ∼7 cm long 1 mm diameter capillary. Importantly, counter-diffusion crystals grew over 1-2 weeks and retained diffraction for up to 2 months. Crystals were harvested from drops or capillaries into a cryoprotectant solution containing 68% mother liquor, 10% PEG3350, 10% ethylene glycol, 10% glycerol, 2% ethanol and cryo-cooled in liquid nitrogen. SeMet-labeled iPLA_2_β was produced in Sf9 cells by using methionine-deficient medium (Expression Systems, Davis, CA) and supplemented with 100mg/L L-selenomethionine 16 h after infection. The I701D mutant, which had 2-3 times greater expression than wild-type, was used for the production of the SeMet protein. Purification and crystallization of the SeMet derivative was the same as for native protein.

### Structure Determination

X-ray diffraction data were collected on GM/CA@APS beamlines 23ID-B and 23ID-D at the Advanced Photon Source, Argonne National Laboratory. Data collection and refinement statistics are shown in Supplementary Table 1. To identify parts of crystals suitable for data collection, more than 400 samples were tested with the raster method using a small (5-20 µm) beam. The best data sets were collected from elongated crystals using the helical method in order to spread the absorbed dose over larger volume of the crystal and thus reduce the radiation damage to the samples^86^. Data were processed and scaled with HKL2000^87^. It was important to use the “Autocorrection” option during scaling. While it reduced the data set completeness and yielded strongly anisotropic data at resolution higher than 4.4 Å (in the highest resolution range (3.95-4.09 Å), 66% of the data had intensity less than 1σ), it also resulted in data with a lower Wilson B factor and significantly more detailed electron density maps.

Data from the SeMet protein crystal were collected at the selenium absorption peak and inflection wavelengths using helical mode and inverse beam geometry with a 30° wedges. Analysis of MAD data at two wavelengths with the Phenix suite^88^ did not yield a solution. SAD data using peak wavelength produced a solution with 7 selenium peaks. A MR replacement solution was obtained using two different protein models, a patatin^62^ and four ARs of an ankyrin-R protein^60^. The structure of patatin was manually trimmed to retain only structural core elements overlapping with CAT domain residues accordingly to the sequence alignment. The Sculptor program within Phenix was used to prepare four ARs from PDB 1N11. The MR solution contained two copies of each domain. Combination of SAD with MR solution resulted in 51 selenium peaks and a high-quality electron density map (Fig. S2a) sufficient for modeling of five additional ARs and several loop regions within the CAT domain. Connectivity of CAT and ANK domains was verified by analysis of all pairs of symmetry-related ANK and CAT domains in the crystal lattice which yielded only one pair with sufficiently short distance. The large number of methionines spread throughout the entire sequence permitted an unambiguous assignment of amino acids. During consecutive steps of structural modeling, combined MR/SAD electron density maps were calculated with one of the domains omitted to avoid model bias. Only one copy of each domain was modeled and the structure was refined using a global NCS function and secondary-structure geometry restriction. After completion of model building, the structure was subsequently refined using 3.95 Å resolution data from the native protein crystal. Simulated annealing composite omit maps were extensively used in model building. Several rounds of Rosetta refinement in Phenix were used for the final model. Phi-psi values of 82% of the residues in the final model are in a favorable region of the Ramachandran plot with 1% in an unfavorable conformation. The latter were in loop regions with poor electron density. Residues 1-80, 95-103, 113-117, 129-145, 405-408, 630-631 and 652-670 were omitted from final model and regions 81-94, 104-112 (numbering in both regions is based on secondary structure prediction) and 409-416 were modeled as alanine residues.

4.6 Å resolution SAD data were collected at selenium peak wavelengths from the protein crystals soaked with 2’MeSe-ATP (Jena Bioscience). Combined MR/SAD analysis revealed a single peak. Several alternative models with different omitted domains or domain fragments were used to avoid model bias. All calculations resulted in an identical position of the selenium peak.

### Fluorescent iPLA_2_β Activity Assay

The continuous activity assay was adapted from a protocol used forsPLA^89^. 1-hexadecanoyl-2-(1-pyrenedecanoyl)-*sn*-glycero-3-phosphocholine (Pyrene-PC, ThermoFisher #H361) (Fig. S5a,b) was dissolved as a 1 mM stock in DMSO. The solution was injected into a glass vial containing assay buffer (25 mM HEPES 7.5, 150 mM NaCl, 10% glycerol) over 1 min with shaking to create the substrate mixture. This method resulted in liposomes averaging 100 nm in diameter as determined by dynamic light scattering. 100 µL of substrate mixture was added to a black 96-well microplate with a non-binding surface (Corning #3650). 0.2% fatty acid free BSA in the buffer acted as an acceptor for the hydrolyzed 1-pyrenedecanoic acid. Proteins were dialyzed against the assay buffer. iPLA_2_β was incubated with different concentrations of CaM with 1 mM CaCl for 15 min. The baseline fluorescence of the substrate was recorded for 3 min at 340 nm excitation / 400 nm emission using the monochromator of a Biotek Synergy 4 plate reader. 10µl of the protein mixture was added to initiate reaction. After a 5 sec mixing step, the fluorescence was read every 30 sec for 1 h or until the signal reached a plateau (Fig. S5c). The linear slope of the first 5 min of the reaction was used as the initial velocity (Fig. S5d, e). The calmodulin inhibition data were fit to the Hill equation using Origin 8.6 software. The velocity in fluorescence units/time was quantified in moles using a curve of the 1-pyrenedecanoic acid product.

### Fluorescence Anisotropy Binding Assays

As calmodulin has no native cysteine residues, a mutant was engineered at Thr34, as described previously^90^, to enable coupling of FAM fluorophore in a site-directed manner. This enabled to measure direct binding of FAM-CaM the using fluorescence anisotropy method. The CaM T34C mutant was created by mutagenesis, confirmed by sequencing, and purified with the same procedure as described for the native protein. The labeled protein was separated from excess FAM with phenyl sepharose in the same procedure as for purification. The concentration of labeled protein was measured at 495nm with a molar extinction coefficient of 68,000 M^-1^ cm^-1^. For the fluorescence binding assay, proteins were dialyzed to the assay buffer (described in activity assay). CaM-FAM (30 nM final concentration) was incubated with a series of iPLA_2_β concentrations obtained by 2-fold serial dilution in a 384 well non-binding plate (Corning #3573) in a total volume of 80µL. After 15 min incubation at 25^°^C, the overall fluorescence intensity and the parallel and perpendicular components were read on a Biotek Synergy 4 with 485 nm excitation and 528 nm emission filters. The fluorescence anisotropy was calculated by the Biotek Gen5 software using the following equation: ***A***=(***F***_∥_−***F***_⊥_)/ (***F***_∥_−***2F***_⊥_), where *F*_∥_ and *F*_⊥_ are the parallel and perpendicular intensities, respectively. Each experiment was conducted in triplicate at least two independent times and values shown are the average ± SEM.

### Analytical Ultracentrifugation

Proteins were extensively dialyzed against AUC buffer (25mM HEPES 7.5, 500mM NaCl, 10% glycerol). Sedimentation velocity studies were performed in a Beckman XL-A analytical ultracentrifuge at 20^°^C and 35,000rpm. The absorbance at 280nm was collected every 4 min for a total of 200 scans. The buffer viscosity and density as calculated by Sednterp (http://www.rasmb.org/sednterp) were 1.04913 ρ and 0.01436 η, respectively. These values were used to fit the data to the Lamm equation in SEDFIT software^91^ using the continuous c(s) distribution model. Graphs were prepared using GUSSI software (UT Southwestern).

## Acknowledgements

We thank Aaron Naatz for help in purifying of *E. coli* expression constructs, Praveen Subramanian for preliminary work on fluorescence assays of PLA_2_ enzymes and all members of the Korolev lab for helpful discussions. We are grateful to Nicola Pozzi, Enrico Di Cera, David Ford, Jane McHowat and William S. Sly for extremely helpful discussions and to Joel Eissenberg for manuscript preparation. GM/CA-CAT beamlines 23-ID-B and 23-ID-D at the Advanced Photon Source, Argonne National Laboratory are funded in whole or in part from the NCI (ACB-12002) and the NIGMS (AGM-12006). This research was supported by the American Heart Association Grant-in-Aid #0665513Z, Center for Advancement of Science in Space (CASIS) grant CASIS-2016-1, NIH/NINDS grant R21NS094854, to S.K. and NIH/NHLBI grant R01HL118639, to R.W.G.

## Author Contributions

K.R.M. and S.K designed, performed, and analyzed experiments, including crystallographic data collection and refinement. C.M.J. and R.W.G. provided purified protein for initial crystallization experiments, plasmid for protein production, and provided expertise in studying the effectors of this enzyme. O.K. cloned multiple protein isoforms in *E. coli* and insect cells, developed protein purification and SeMet labeling protocols and contributed to crystallization and activity measurements. I.M. assisted with cloning of mutants, cell culture, protein purification, activity and AUC assays. R.S. collected data set from native protein crystal and helped with data collection strategies for selenium protein derivatives. K.R.M. and S.K wrote the paper with input from all authors.

## Accession codes and data availability

Atomic Coordinates and structure factors for the iPLA_2_β structure have been deposited in the Protein Data Bank under accession code PDB ID 6AUN. All reagents and relevant data are available from the authors upon request.

## Supplementary Data

**Table S1.**
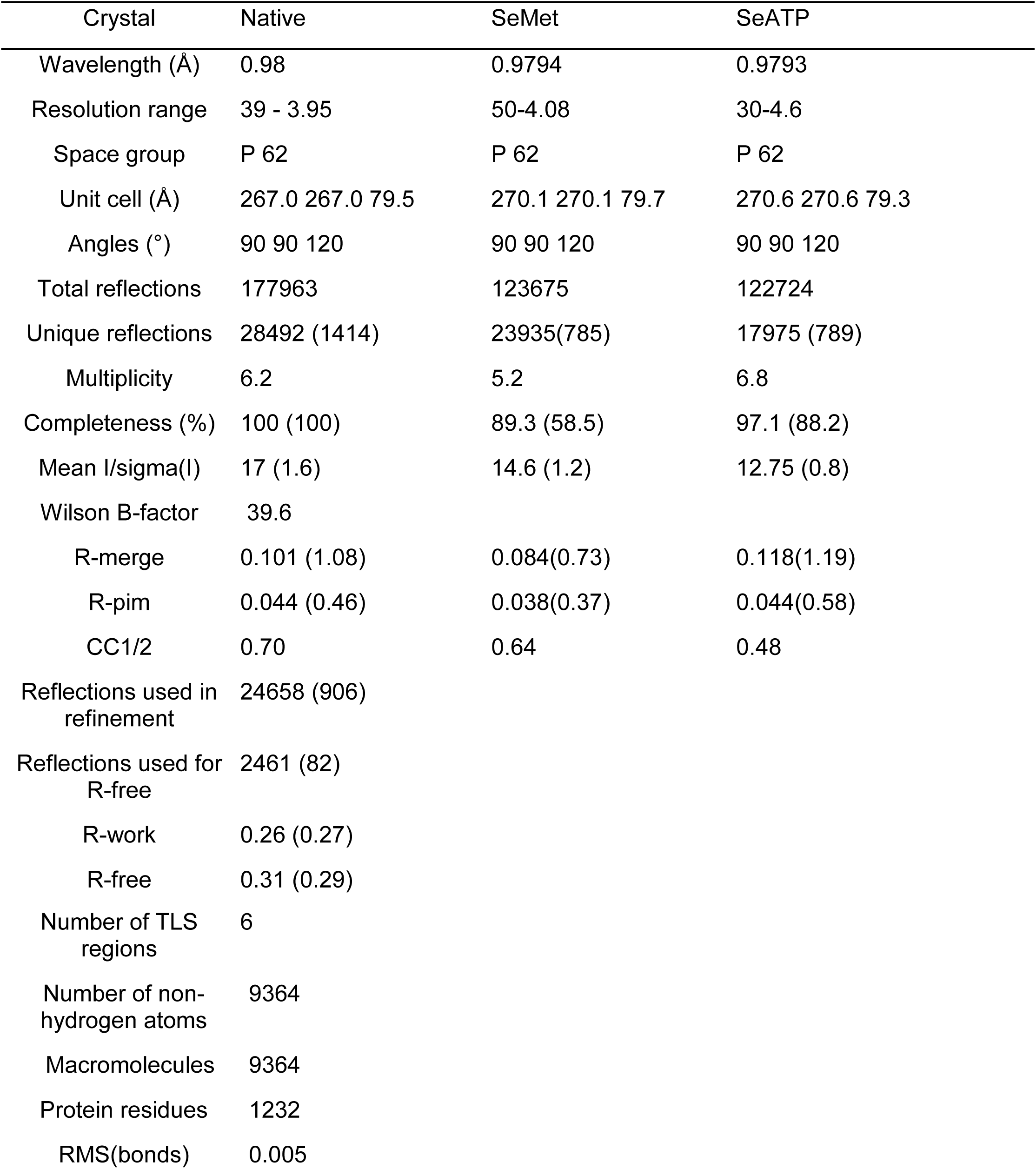
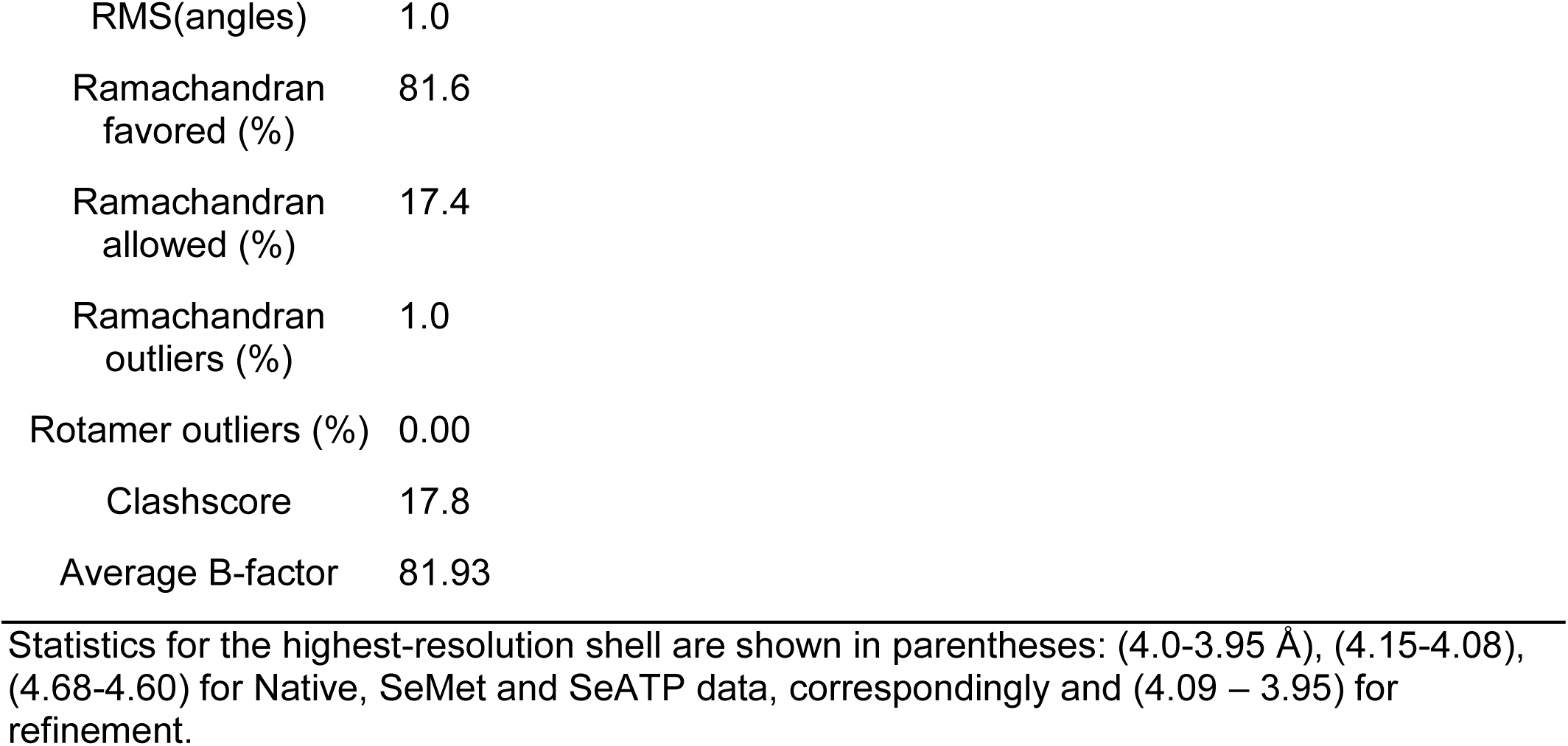
Data collection and Refinement statistics

| Crystal | Native | SeMet | SeATP |
| --- | --- | --- | --- |
| Wavelength (Å) | 0.98 | 0.9794 | 0.9793 |
| Resolution range | 39 - 3.95 | 50-4.08 | 30-4.6 |
| Space group | P 62 | P 62 | P 62 |
| Unit cell (Å) | 267.0 267.0 79.5 | 270.1 270.1 79.7 | 270.6 270.6 79.3 |
| Angles (°) | 90 90 120 | 90 90 120 | 90 90 120 |
| Total reflections | 177963 | 123675 | 122724 |
| Unique reflections | 28492 (1414) | 23935(785) | 17975 (789) |
| Multiplicity | 6.2 | 5.2 | 6.8 |
| Completeness (%) | 100 (100) | 89.3 (58.5) | 97.1 (88.2) |
| Mean I/sigma(I) | 17 (1.6) | 14.6 (1.2) | 12.75 (0.8) |
| Wilson B-factor | 39.6 |  |  |
| R-merge | 0.101 (1.08) | 0.084(0.73) | 0.118(1.19) |
| R-pim | 0.044 (0.46) | 0.038(0.37) | 0.044(0.58) |
| CC1/2 | 0.70 | 0.64 | 0.48 |
| Reflections used in refinement | 24658 (906) |  |  |
| Reflections used for R-free | 2461 (82) |  |  |
| R-work | 0.26 (0.27) |  |  |
| R-free | 0.31 (0.29) |  |  |
| Number of TLS regions | 6 |  |  |
| Number of non-hydrogen atoms | 9364 |  |  |
| Macromolecules | 9364 |  |  |
| Protein residues | 1232 |  |  |
| RMS(bonds) | 0.005 |  |  |

|  |  |
| --- | --- |
| RMS(angles) | 1.0 |
| Ramachandran<br>favored (%) | 81.6 |
| Ramachandran<br>allowed (%) | 17.4 |
| Ramachandran<br>outliers (%) | 1.0 |
| Rotamer outliers (%) | 0.00 |
| Clashscore | 17.8 |
| Average B-factor | 81.93 |

Statistics for the highest-resolution shell are shown in parentheses: (4.0-3.95 Å), (4.15-4.08), (4.68-4.60) for Native, SeMet and SeATP data, correspondingly and (4.09 – 3.95) for refinement.

**Figure S1.**
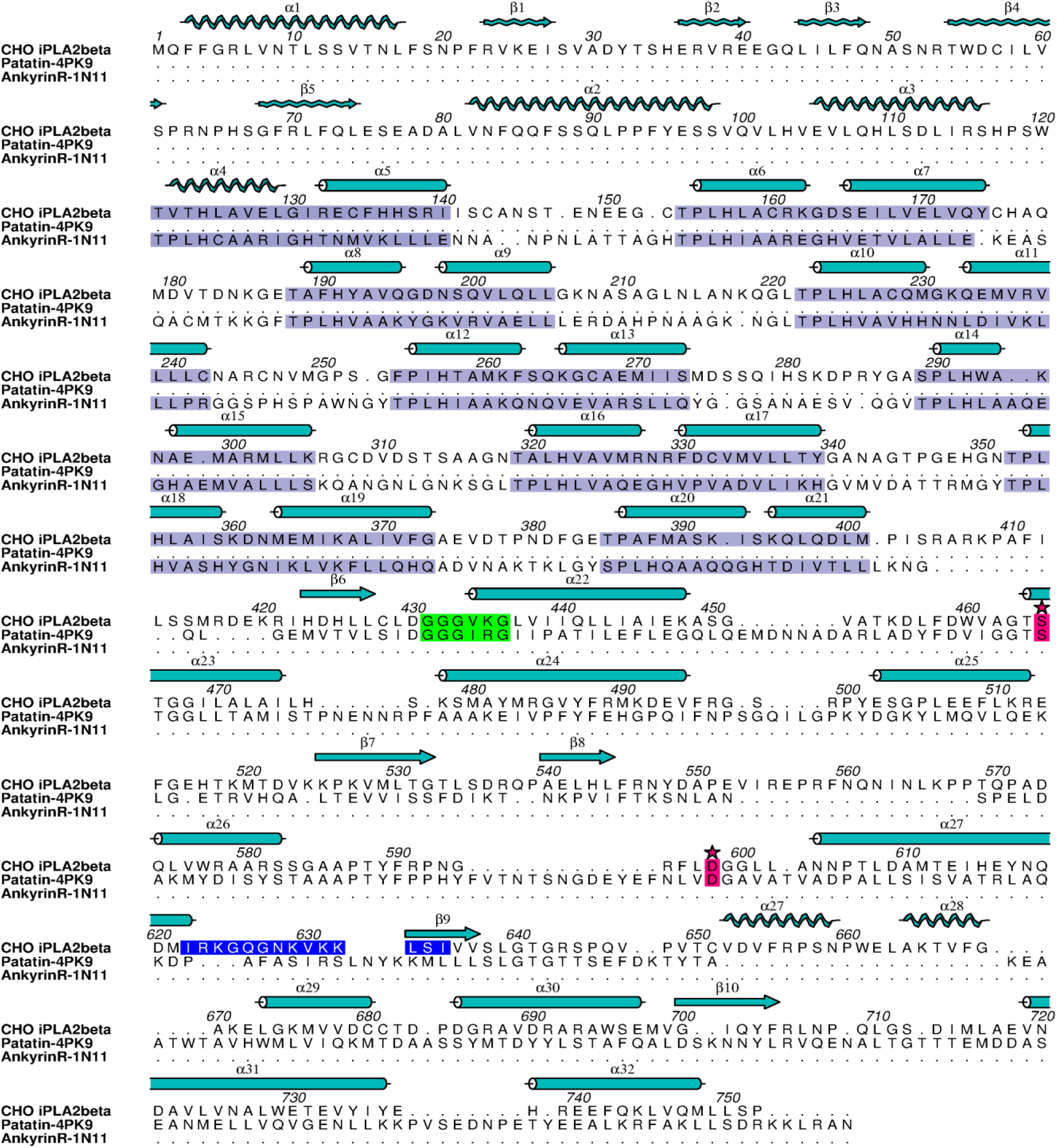
Structure-based alignment of the iPLA_2_β amino acid sequence with patatin (PDB: 4PK9) and the eight ankyrin repeats of ankyrin-R (PDB: 1N11) using TM-align. The ARs are highlighted in light purple, the active site residues starred in magenta, the oxyanion hole in green, and the 1-9-14 CaM-binding motif in blue. Helices and sheets from the structure are depicted with cylinders and straight arrows, respectively, and predicted secondary structure elements in the N-terminus and membrane interacting regions are shown with wavy lines and arrows.

**Figure S2.**
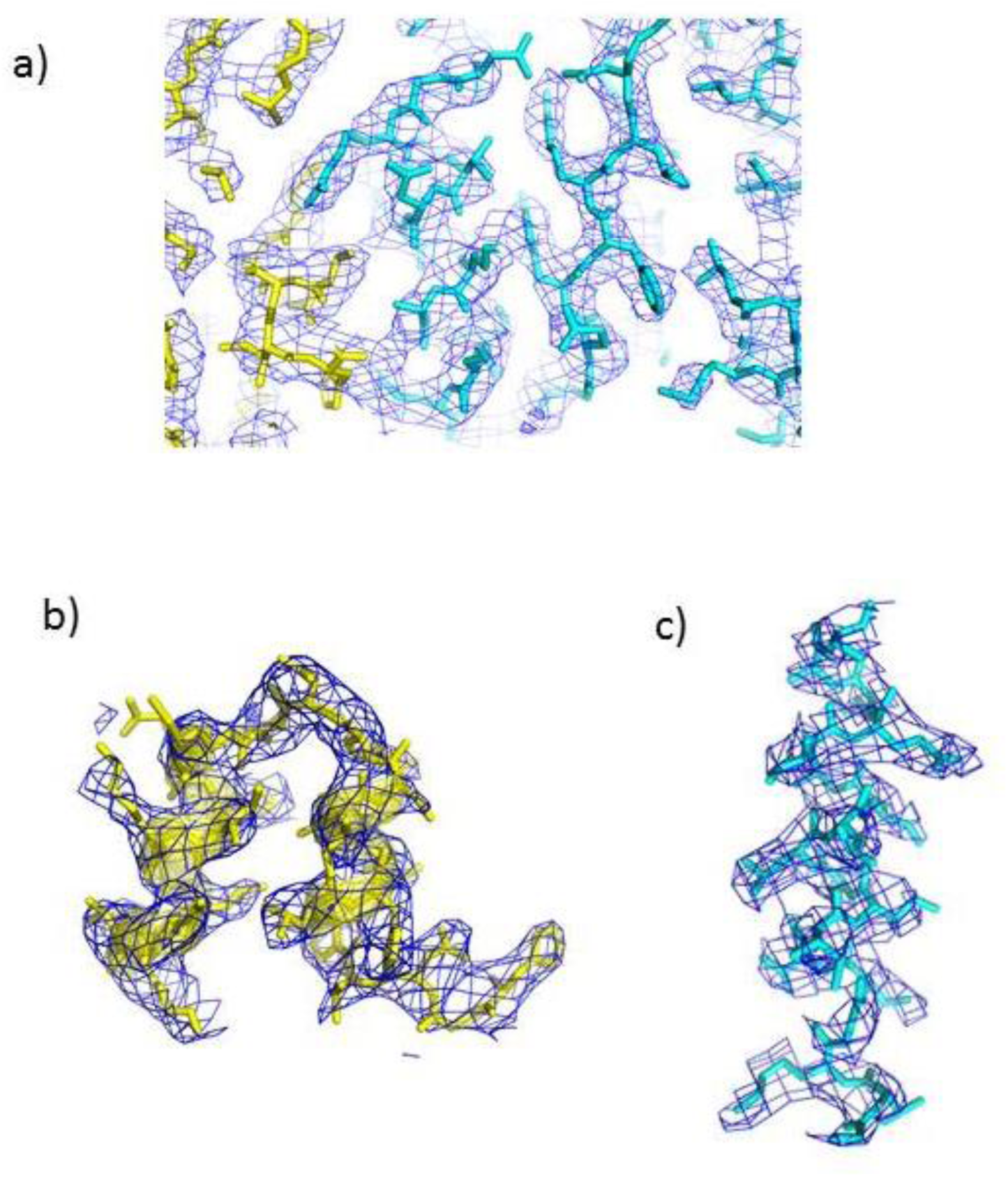
iPLA_2_β structure determination by MR/SAD. 2Fo-Fc **e**lectron density maps around several regions of the protein, including **a)** the dimerization interface, **b)** AR_9_ and **c)** N-terminal α-helix contoured at 1.5σ.

**Figure S3.**
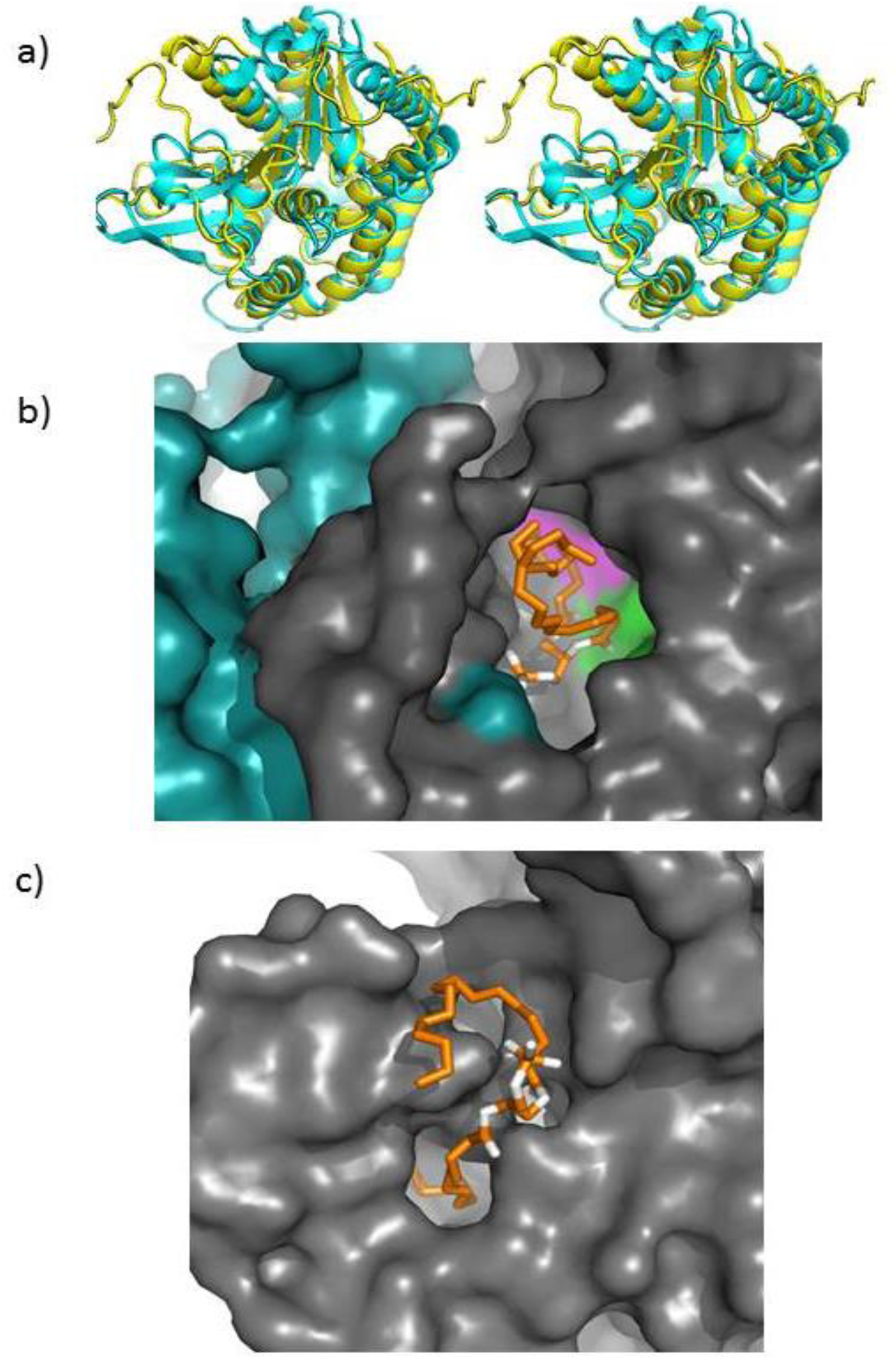
Comparison of patatin and iPLA_2_β CAT domains. **a)** Stereo view of the superposition of the iPLA_2_β catalytic domain shown in yellow with patatin (PDB ID:4PK9), shown in cyan; **b)** An open conformation of the active site of iPLA_2_β. The surface representation of the iPLA_2_β active site region (the view from the membrane) is shown in grey for monomer A and in cyan for monomer B. Active site residues are highlighted in green and magenta. To illustrate the open conformation and accessibility of the active site, the 1,2-dipalmitoyl-3-sn-phosphatidylethanolamine molecule shown with stick representation in orange and white was inserted using the Dock program. **c)** Surface representation of the membrane-interacting region of patatin in similar orientation. Catalytic residues are completely inaccessible and only one fatty acid chain can be fitted into narrow cavity leading to the active site.

**Figure S4.**
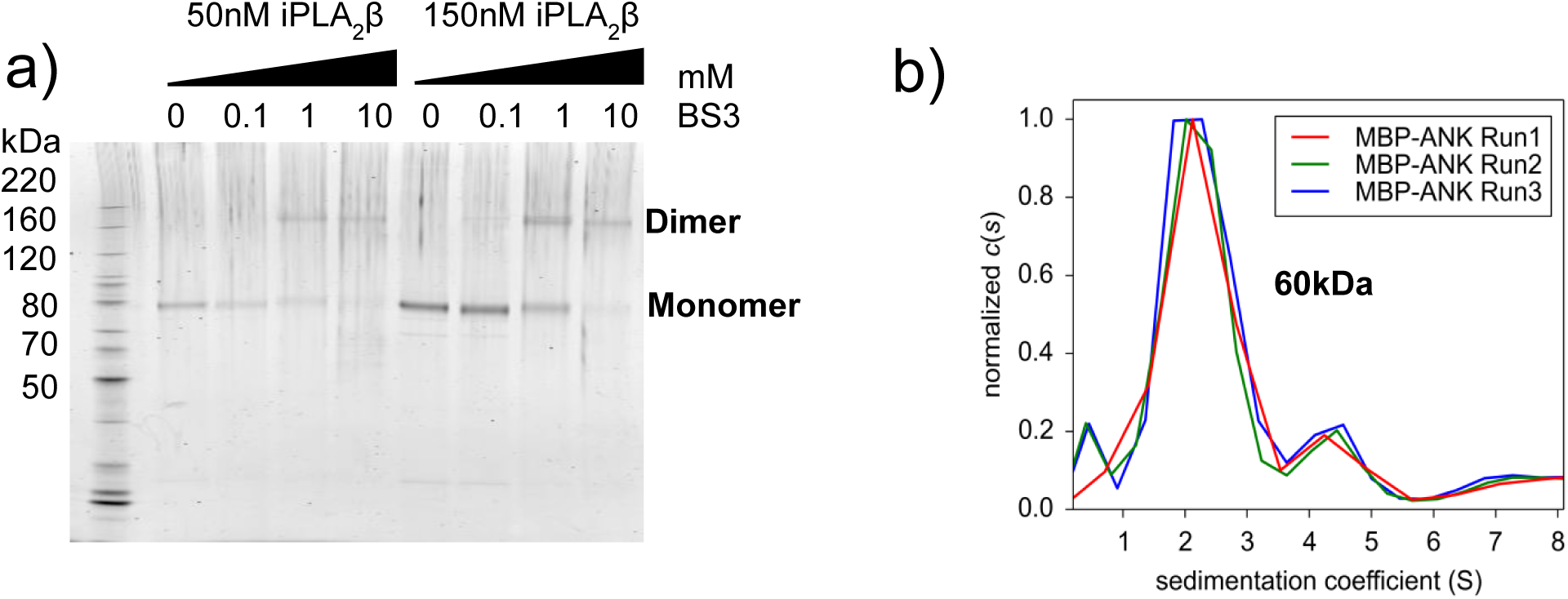
iPLA_2_β dimerization in solution. **a)** Dimerization of iPLA_2_β revealed by crosslinking experiments. The protein was treated with increasing concentrations of BS3 amine-reactive crosslinking agent (concentration in mM is shown above the gel) for 1 hour. The reaction was quenched by Tris-glycine loading buffer, proteins were separated by SDS PAGE, stained with Krypton Fluorescent Protein Stain, and visualized at 523 nm excitation and 580 nm emission on a Typhoon imager. **b)** C(s) distribution of purified MBP-tagged ANK domain (177-389) from three independent sedimentation velocity experiments. The estimated molecular weight corresponds to a monomer of the MBP-ANK (average MW of peaks is 60kDa versus theoretical 63 kDa).

**Figure S5.**
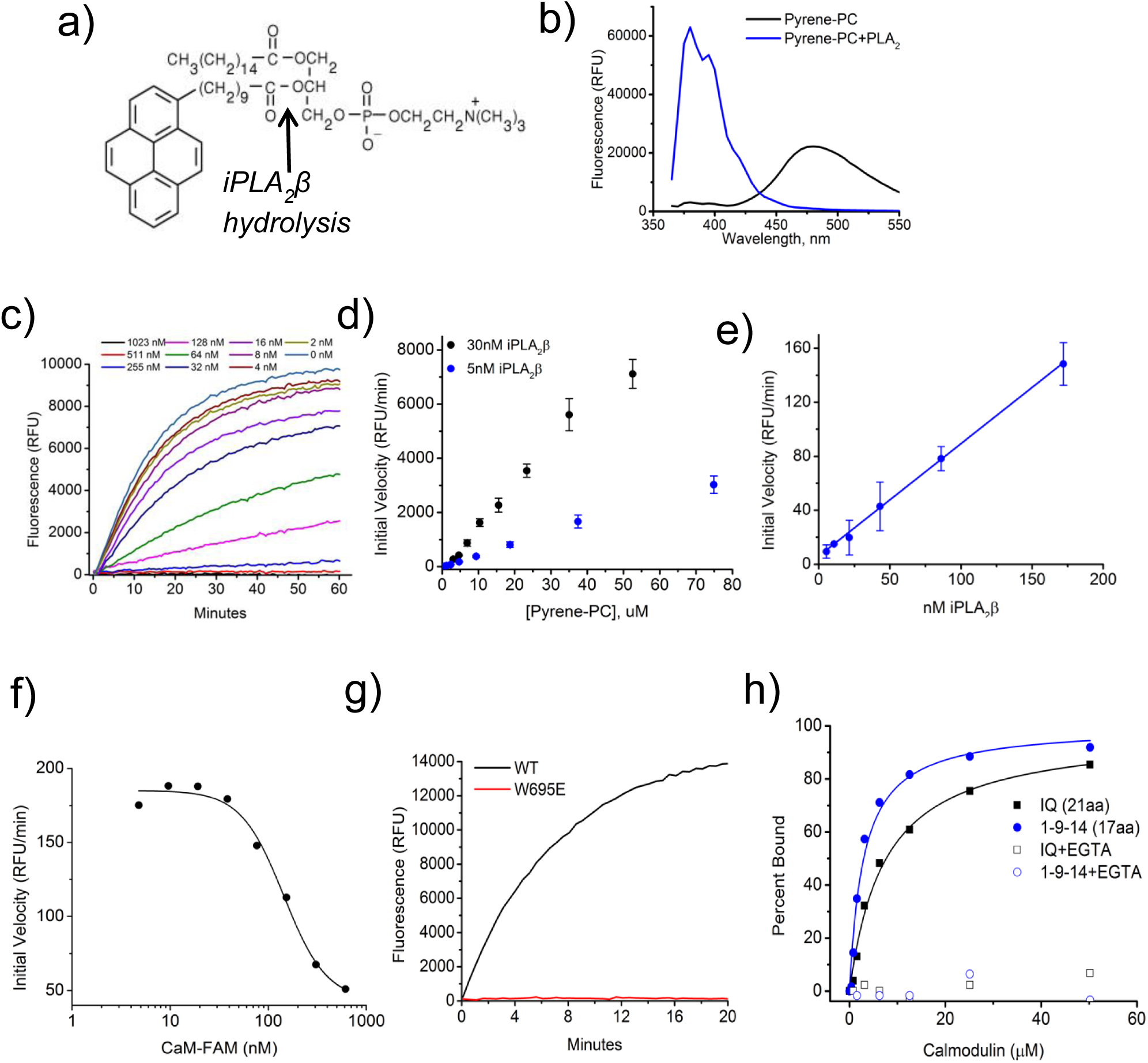
Enzymatic activity and inhibition measured with pyrene-PC substrate. **a)** Structure of pyrene-PC. **b)** Fluorescence spectra at 342 nm excitation of the substrate before and after cleavage. **c)** Real time measurements of pyrene-PC cleavage by iPLA_2_β in the presence of different concentrations of CaM. RFU – relative fluorescence unit. The initial slope of these data was used in preparing Fig 4c. **d)** Plots of initial velocity vs. substrate concentration at two different concentrations of enzyme. **e)** Validation of the assay demonstrating the linear relationship between initial velocity and the enzyme concentration. Error bars represent average ± s.e.m of assays performed in triplicate. **f)** Inhibition of iPLA_2_β by FAM-CaM. **g)** Activity of the wt protein and W695E mutant. **h**) Binding of IQ and 1-9-14 FAM-labeled synthetic peptides to CaM measured by fluorescence anisotropy. Data is representative of three independent experiments.

**Figure S6.**
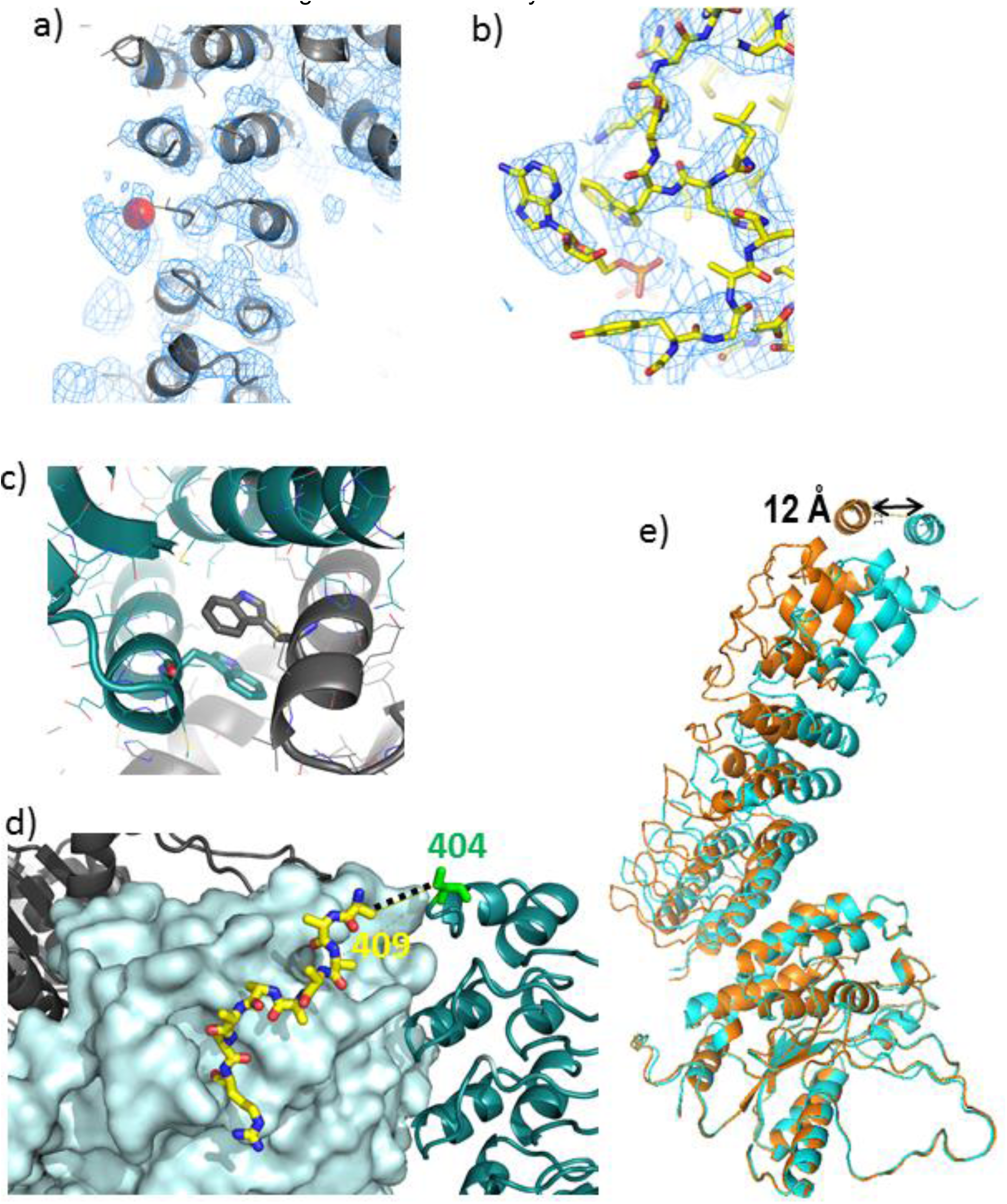
**a)** Position of the Se peak (red sphere) found from the Se-ATP soaked crystal and shown with the experimental electron density map. **b**) Fitting of ATP into the electron density map of the SeMet data. **c)** Interactions of Trp695 shown in stick representation in a dimerization interface. One molecule is shown in grey and the other in dark cyan. **d)** Conformation of partially resolved loop, shown in yellow, red and blue stick representation, connecting CAT and ANK domain. CAT domain is shown in surface representation and ANK in cartoon representation. **e**) Movement of ANK upon superimposition of CAT domains. One molecule is colored in orange and the other in cyan.

